# A human spiking computational model to explore sound localization

**DOI:** 10.1101/2025.09.18.677043

**Authors:** Francesco De Santis, Paolo Marzolo, Alessandra Pedrocchi, Alberto Antonietti

## Abstract

Sound localization relies on precise processing of binaural cues in medial (MSO) and lateral superior olive (LSO). However, key questions remain on how these two nuclei perform their specific computations depending on sound frequency, the relative contributions of interaural time differences (ITDs) and interaural level differences (ILDs), as well as the role of inhibitory timings. Experimental studies have struggled to address these issues because of technical challenges and a lack of methodological consistency. Here, we present a comprehensive computational model of auditory peripheral and brainstem neural populations to investigate how ITDs and ILDs are encoded by LSO and MSO. We developed a spiking neural network with realistic tonotopic organization and biologically consistent synaptic connections. We tested responses to pure tones and white noise from different locations under three cue conditions: human-recorded head-related transfer functions, isolated ITDs, and isolated ILDs. LSO neurons showed realistic ipsilateral-preferring responses across different stimuli, with cue dependency varying by tone frequency, while white noise responses were driven mostly by ILDs. MSO responses showed heterogeneous tuning, with contralateral preference for low-frequency tones, which got lost for higher-frequency ones, and white noise response driven by ITDs. We further validated two experimental findings: (1) removing MSO inhibition abolished contralateral tuning, and (2) varying the timing between excitation and inhibition produced large shifts in tuning, highlighting the importance of synaptic timing for ITD coding. This model serves as an *in silico* testbed for auditory research, offering new insights into the functioning of human spatial hearing.

## 1. Introduction

**S**OUND localization is a fundamental function of the human auditory system, enabling individuals to identify and react to environmental sounds. Unlike other sensory systems, such as vision or somatosensory, the auditory system does not present spatial organization at the receptor level. Instead, inner hair cells (IHCs) transducing sound are arranged along the cochlea according to their sound frequency tuning, which is defined as their characteristic frequency (CF). Therefore, our brain must construct acoustic spatiality through the processing of the two sound inputs arriving at our ears, especially by leveraging hidden spatial cues. These spatial cues can be split into two classes: binaural and spectral cues. Binaural cues are interaural time differences (ITDs) and interaural level differences (ILDs), which respectively consist of the difference in arrival time and intensity of sound between the two ears. Spectral cues refer to the filtering effects introduced by the head and pinnae; they are also called monaural, since they can be detected in single ear response. In humans, psychoacoustic studies prove that binaural cues alone are sufficient to discriminate azimuth angles in the frontal plane, ranging from −90° to +90°, which correspond to the extreme left and right sides of the head [1], [2]. Spectral cues, instead, solve front-back ambiguities and angle recognition along the vertical plane (i.e., elevation angles) [3], [4]. The Head-Related Transfer Function (HRTF) mathematically represents the combined effect of binaural and monaural cues, including every acoustic transformation that sound undergoes on its path from source to ear. As such, they depend on the anatomy of each subject.

### A. Anatomy and physiology of the brainstem

Once transduced to neural spike trains by IHCs, all spatial cues are conveyed by the input spike trains of the auditory nerve fibers (ANFs). Therefore, understanding ANF behavior is essential to examining how brainstem auditory circuits extract and analyze spatial cues. Neurons involved in spatial hearing show two properties introduced by ANFs and preserved by higher populations: tonotopic organization and phase locking. Tonotopic organization originates from the graded mechanical properties of the cochlear basilar membrane and the CF of IHCs along its length: each IHC, and therefore each ANF it contacts, is most sensitive to a specific CF. This frequency-based spatial arrangement allows the brain to analyze sound content based on spectral composition. Phase locking is the ability of ANFs to fire action potentials at a particular phase of a sound wave, enabling precise temporal encoding of acoustic signals [5], [6]. This precise timing is fundamental for sound processing and, in the context of our investigation, it is critical for processing ITDs. Spherical and globular bushy cells (SBCs and GBCs) in the anteroventral cochlear nuclei receive convergent ANF inputs and sharpen phase-locked timing even further, refining the temporal precision needed for accurate ITD detection [5], [7]. From bushy cells, precisely timed excitatory outputs project both ipsilaterally and contralaterally through a network of excitatory and inhibitory pathways, involving as intermediate synaptic stages the glycinergic cells of medial and lateral nuclei of the trapezoidal body (MNTB and LNTB) and as final target the medial and lateral olives in the superior olivary complex (MSO and LSO). Principal neurons in the LSO receive excitatory input from ipsilateral spherical bushy cells (SBCs) and inhibitory input from contralateral MNTB neurons, forming an excitation-inhibition circuit for sound localization. MSO neurons have two additional inputs, consisting of contralateral excitation from SBCs and ipsilateral inhibition from LNTB cells. Depending on how relevant inhibitory inputs were considered in coding the location of sound, the dynamics of MSO neurons have been described as excitation-excitation [8]–[12], inhibition-excitation-excitation [13]–[15] or inhibition-excitation-inhibition-excitation [16], [17]. When presented with sounds coming from different frontal azimuth angles (between −90° and +90°), the LSO exhibits a sigmoid-shaped rate response curve, showing a preference for ipsilateral sounds. Firing rates of left and right LSOs converge at 0° and progressively diverge for more lateral sound sources [18]. In contrast, MSO neurons show contralateral preference [14], [19], [20], with firing rate curves shown to be sigmoid-like (reaching a plateau for contralateral angles between 0° and 90°) or peaked (reaching a maximum at a specific contralateral angle after 0° before decreasing again toward 90°) [5].

### B. State of the art

Despite more than a century of neuroscientific research, key questions about how we localize sound remain unanswered. For example, the dichotomy between ITDs and ILDs as a function of tone frequency, and their respective processing in the MSO versus the LSO, is still debated. The prevailing explanation is still the duplex theory, which states that the auditory system relies on ITDs to localize low-frequency sounds and on ILDs for high-frequency sounds [2]. This theory is supported by both physiological and psychophysical evidence: in humans, IHCs and consequently ANFs lose the ability to phase-lock to sound waveforms at frequencies above approximately 3 kHz due to inherent biological constraints [6]. This degradation of phase-locking renders ITD cues ineffective at high frequencies, where ILDs (generated by the acoustic shadow created by the head) become instead the dominant spatial cue. Numerous psychoacoustical studies with pure tone stimuli have corroborated this frequency-based division of labor between ITDs and ILDs, also showing how for tones with mid-to-high frequency (approximately 2 to 4 kHz) humans exhibited particularly poor localization acuity, since these frequencies are too high for effective ITD processing (due to reduced phase-locking), yet too low to produce strong ILDs [21]. However, several critical issues remain open: first, the duplex theory was mainly validated using pure tones [21], while complex stimuli, including white noise or speech, are still under investigation [5]. Second, the physiological underpinnings of ILD and ITD processing within the brainstem nuclei in humans, especially the extent to which the MSO and LSO are exclusively specialized for one cue type, remain unclear. Recent findings suggest that the relationship between ILDs, ITDs, and their respective nuclei may not be as univocal and functionally segregated as once thought [18], [22]. Nevertheless, traditionally the MSO has always been considered the main processing site of ITDs. This has sown the seeds for numerous theories on the dynamics of its neurons: ITDs are on the order of microseconds (depending on the frequency of the sound and the size of the head), whereas a neural action potential has a duration of 1-2 ms, so the mechanism behind their detection and discrimination is not trivial. For almost a century, hypotheses have held true to the Jeffress model [8], which suggested coincidence detection by means of excitatory-excitatory dynamics. Nevertheless, in the last twenty years, many studies have challenged this simplistic view, claiming that inhibitory inputs could play a crucial role in shaping the temporal precision of MSO neurons and consequently ITDs discrimination [13]–[17]. Pecka and colleagues [14] provided physiological and pharmacological evidence from *in vivo* extracellular MSO recordings in anesthetized gerbils, in which reversible blockade of synaptic inhibition by glycine antagonist strychnine increased firing rates and significantly shifted ITD sensitivity of MSO neurons. Myoga and colleagues [17] applied temporal-precise stimuli using conductance-clamp to adult gerbil brain slices, demonstrating that the combined effects of the two inhibitory inputs can dynamically shift the timing of peak in the rate curve, depending on their relative arrival times. Specifically, by artificially varying the delays between each inhibitory input and its corresponding ipsilateral excitation (Δ*T*^*inhi*^ for the ipsilateral pair and Δ*T*^*inhc*^ for the contralateral pair) Myoga was able to modulate the timing of optimal coincidence detection. Despite this evidence, many other studies [9]–[12] have reported opposite results, denying the role of inhibition in shaping MSO rate response and ITD’s sensitivity. Moreover, Myoga’s results about the effect of Δ*T*^*inhi*^ and Δ*T*^*inhc*^ on the timing of peak firing in MSO neurons are based on single-cell analyses. To date, no evidence demonstrates that the same mechanism operates across the entire sound localization network. A key reason why so many questions in sound localization research remain unsolved is the combination of technical challenges and a lack of methodological homogeneity between studies. Recording from the tiny, fiber-dense MSO and LSO buried deep in the brainstem demands highly precise stereotaxic access and often causes tissue disruption that can compromise signal stability [5], [23]. At the same time, experimental protocols vary widely: investigators use different animal models (gerbils [9]–[15], [17], chinchillas [24], guinea pigs [19], [25], bats [26], cats [7], [18], [20], [22]), employ diverse stimuli (pure tones [13], [26], tone bursts [27] wideband noise [5], [19], [22], clicks [28]), introduce binaural cues through realistic HRTFs [29], isolated ITD [14], [17], [19], [20] or ILD presentation [26], [27], or binaural beats [5], [11], and even target different nuclei (inferior colliculus in place of MSO [5]). Finally, heterogeneity in cell selection (spanning a range of CFs) complicates comparisons. Together, these technical and procedural inconsistencies make it difficult to integrate findings and advance a unified understanding of spatial cues processing.

### C. Computational modeling of sound localization

This particular context is a fertile ground for the development of computational models, which offer systematic manipulation of variables (sound stimuli, cue applications, and network parameters) and easy access to all simulated populations at each time instant. Several spiking neural network (SNN) models for sound localization have recently been proposed, each offering valuable insights yet addressing only part of the story. A leaky integrate-and-fire network modeling MNTB and LSO neurons and trained with cat HRTFs excelled at high-frequency localization but ignored ITD cues, failing at low-frequency localization [30]. A Hodgkin-Huxley-based model simulated ANFs, GBCs, and MSO, but omitted the other involved populations and required a separate artificial neural network to decode timing [31]. Another approach combined MSO and LSO populations feeding into an inferior colliculus (IC), demonstrating that ILDs improve localization above 1 kHz, yet computed IC responses via Bayesian inference rather than spiking dynamics [32]. A minimal GBC coincidence-detector captured biological variability efficiently but stopped short of downstream MSO/LSO processing [33]. Finally, a complete *in silico* circuit covering ANFs to MSO and LSO populations was proposed in [34] but still relied on approximated input spike trains in place of real recorded sounds.

In this work, we introduce a comprehensive, realistic, human-specific network that replicates all relevant stages, from cochlear encoding to MSO and LSO populations, driven by realistic sounds filtered by authentic human HRTFs and neuronal populations parameterized with human (or related mammalian) data. With our results, we aim to clarify the role of ITD and ILD in MSO and LSO activity when stimulated by different input sounds. We therefore shed light on the encoding of sub-millisecond ITDs in the MSO by endorsing the claims on the role of well-timed inhibition in shaping its rate-response. Finally, we offer this model as a potential testbed for replicating experimental setups and advancing research in sound localization.

## II. Methods

The implemented computational model was developed with the main objective of maximizing its plausibility. This concerned both the filtering transformation of sound from the source in the environment to the subject’s ear canal and the receptor transduction of sound waves into spiking activity, as well as the consequent processing performed by higher-level neural populations. As such, the requirements of the two stages varied substantially, and different strategies were adopted. The modeling of the peripheral section, from sound to the spiking output of ANFs, was implemented using Brian2Hears [35], an extension to Brian 2 simulator [37]. The modeling of the brainstem nuclei involved in sound localization was developed using the NEST simulator framework [36].

### A. Peripheral Modeling

#### 1) Sounds

To comprehensively evaluate how our network processes sound frequencies, we selected four different acoustic stimuli: pure tones at 0.1 kHz, 1 kHz, and 10 kHz, representing low, mid, and high frequency ranges, respectively, along with white noise, capable of simultaneously activating all IHCs along the cochlea. All four selected sounds were implemented as built-in Brian2Hears objects, each lasting 1 second and presented at 70 dB intensity. The Brian2Hears framework also enables the generation of more sophisticated stimuli used in auditory research, such as click trains with varying intervals or binaural beats for studying phase-locking and binaural processing, though these complex sounds were not included in the current investigation.

#### 2) Cue Application

In order to address the different contributions of the two main auditory cues associated with sound localization, we tested three different configurations, HRTF, *ITDonly* and *ILDonly*:

- ***HRTF:*** Sound signals were filtered through realistic, subject-recorded HRTFs from the IRCAM LISTEN database [38] to generate binaural inputs for each ear. The HRTFs were acquired through a single-microphone, single-speaker recording setup with a metallic U-shaped supporting structure constraining spatial resolution to 15° azimuthal increments, providing 13 distinct positions. To ensure consistency across experimental conditions, this 15° resolution limit was maintained for all three spatial configuration methods. To ensure robustness, we validated our results for the four sounds studied with 5 HRTFs recorded from different subjects (see *Supplementary*).
- ***ITDonly:*** To isolate the effect of ITDs, we implemented a simplified “headless” model that computes them artificially, without modeling the complex filtering effects of the head, ears, and torso. The *HeadlessDatabase* generates ITDs for 13 equally spaced angles between −90° and +90° using the formula

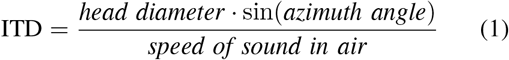 where the diameter represents the distance between the ears (set to give a maximum ITD of 650 *µs*). For each angle, the system creates simple impulse responses that delay the sound to one ear relative to the other. If the ITD is positive, the left ear receives the undelayed sound while the right ear is delayed, and vice versa for negative ITDs. This approach isolates the timing cues for spatial hearing by implementing pure time delays without any of the spectral filtering that would normally occur due to head-related acoustic effects.
- ***ILDonly:*** We developed an interaural level difference (ILD) model that creates artificial amplitude differences between the ears without any temporal delays. We adopted two approaches depending on the sound type: for tonal sounds, we applied a frequency-dependent shadowing model based on acoustic theory, calculating the head shadowing effect using the parameter and limiting the maximum shadowing to 20 dB [4]:

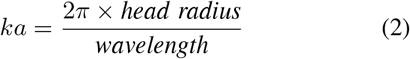

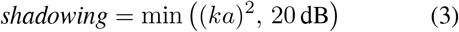

with the final ILD computed as

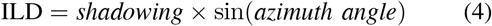

to provide directional sensitivity. For non-tonal sounds (i.e., white noise), we used a simplified linear interpolation that scales linearly with angle, providing up to ±15 dB difference at ±90°. This level difference was chosen according to the realistic ILD value imposed by HRTF functions from the IRCAM LISTEN database corresponding to ±90°, recorded when applied to white noise sound. Finally, the spatial effect was implemented by attenuating the sound level in one ear - subtracting the ILD at the left ear level for rightward sources (positive angles) and at the right ear level for leftward sources (negative angles) - while keeping the original sound timing intact in both channels, thus isolating the amplitude-based spatial cues from timing-based ones.

#### 3) Ear

To model the physical transformations occurring to sounds once entered in our ear canal up to the cochlear basilar membrane, we employed the Tan-Carney model [39]. Unlike simpler models that use a gammatone filterbank followed by rectification, the Tan-Carney model integrates more physiologically accurate mechanisms at multiple stages. First, the sound is filtered through a bandpass filter mimicking middle ear amplification. Then, the model cochlear stage employs a two-path architecture: a time-varying bandpass filter processes the signal path while a nonlinear control path accounts for basilar membrane compression, a crucial nonlinearity in cochlear mechanics. To implement the time-varying bandpass filter, we spaced the center frequencies according to the ERB scale and used 3500 channels, matching the estimated number of inner hair cells (IHCs) in the human cochlea [40], ensuring a biologically realistic tonotopic organization. The inner hair cell (IHC) and ANF synapse were instead replicated using the Zhang et al. model [41], which includes a saturating low-pass filter for the IHC and a three-store vesicle release mechanism to simulate synaptic adaptation, refractoriness, and neurotransmitter depletion dynamics. This synapse model reproduces the temporal and adaptive properties of high spontaneous rate fibers without requiring added noise. We set up 10 Zhang synapses for each IHCs, in order to have a realistic number of ANFs, i.e., 35000 (in humans, type I afferent ANFs are estimated to be 30000-40000 [5], [6]). While originally fitted to cat data, especially low-frequency IHCs, the model remains suitable for human auditory simulations, since human hearing does not extend beyond 20 kHz.

### B. Neural Processing

Once determined by the peripheral model, the ANF spiking patterns are saved into a binary file to avoid re-generation for every run. Then, spike generators are created using the ANF spiking patterns, which will be the inputs for the neural processing section. For the modeling of this phase, we used NEST Simulator [36]. Like Brian 2, NEST allows the modeling of Spiking Neural Networks, enabling experiments to be performed *in silico*. We interacted with the NEST simulator using PyNEST. This Python interface serves as an API layer translating Python requests into code that finally controls the Nest Kernel, written in C++. One significant advantage of NEST was the availability of built-in thread parallelism, which enabled a substantial speedup. We used NEST version 3.3 [**?**] and the Python interpreter version 3.8.10.

#### 1) Network Structure

The first layer of the network consists of an array of NEST *spike generator*s, emulating the human ANFs. To implement the second layer, i.e., SBCs and GBCs of both the cochlear nuclei, we started from convergence data available in literature: SBCs in cats receive 1 to 4 ANF terminals on their cell bodies (we chose to keep 4) [5], [42] and GBCs receive about 20 for each cell (we chose to keep 20) [5], [43]. From this data, we derived neuron counts for the two populations. From this point to the higher nuclei (MNTB and LNTB, then MSO and LSO), little information is available regarding the exact convergence and neuron counts of the populations involved. In our model, GBC neuron counts are the baseline; thus, we obtained a convergence of 5:1 for direct excitatory synapses between SBCs and MSO/LSO PCs and 1:1 for all other connections.

All previously described populations were modeled with the NEST *iaf cond alpha* model, an implementation of a spiking neuron using integrate-and-fire dynamics with alpha-shaped conductance-based synapses. The basic equation describing the dynamics of the neurons’ membrane potential, *V*, can be summarized as

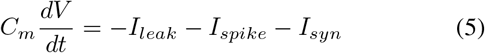

where *C*_*m*_ is the membrane capacitance, *I*_*leak*_ is the current due to the passive leak of the membrane, *I*_*spike*_ is a current describing the spiking mechanism of the neurons, and *I*_*syn*_ is a current describing the effect of synaptic input on the neuron. All cells involved in the auditory pathway have exceptionally fast membrane time constants *τ*_*m*_ [44] which, by definition, describes how the membrane behaves when its only conducting pathway is the passive (leak) conductance. In *iaf cond alpha, I*_leak_ = *g*_*L*_ (*V*_*m*_ *E*_*L*_), where *g*_*L*_ is the leak conductance and *E*_*L*_ the leak reversal potential. Thus, the membrane time constant results equal to *τ*_*m*_ = *C*_*m*_*/g*_*L*_. To shorten *τ*_*m*_ to physiological values, we set a low *C*_*m*_ and a high *g*_*L*_, since a high leak conductance simulates the fast ion channels present throughout the auditory pathway. Table II collects the membrane capacitance and leak conductance.

**TABLE I.** Numerosity (N), Model, and Convergence per neuron population.

| Cell Type | N | Model | Convergence | Ref |
| --- | --- | --- | --- | --- |
| ANFs | 35k | spike_generator | 4:1 → SBC<br>20:1 → GBC | [5], [6] |
| SBCs | 8.75k | iaf_cond_alpha | 5:1 → LSO<br>5:1 → MSO | [5], [42] |
| GBCs | 1.75k | iaf_cond_alpha | 1:1 → LNTB<br>1:1 → MNTB | [5], [43] |
| LNTB | 1.75k | iaf_cond_alpha | 1:1 → MSO | - |
| MNTB | 1.75k | iaf_cond_alpha | 1:1 → LSO<br>1:1 → MSO | - |
| MSO | 1.75k | iaf_cond_alpha | - | - |
| LSO | 1.75k | iaf_cond_alpha | - | - |

**TABLE II.** Membrane capacitance (*C*_*m*_), leak conductance (*g*_*L*_), and time constant (*τ*_*m*_ = *C*_*m*_*/g*_*L*_) for the different neuron populations.

| Population | $C_m$ (pF) | $g_L$ (nS) | $\tau_m$ (ms) |
| --- | --- | --- | --- |
| SBC | 15 | 40 | 0.375 |
| GBC | 15 | 25 | 0.600 |
| MNTBC | 15 | 25 | 0.600 |
| LNTBC | 15 | 25 | 0.600 |
| MSO | 20 | 80 | 0.250 |
| LSO | 30 | 20 | 1.500 |

All other model parameters were kept at default values. Another fundamental class of parameters in our network is synaptic delays. Delays were particularly crucial to study the precise temporal processing in LSO and MSO. In particular, in the MSO, we needed to test the effectiveness of inhibitory ipsilateral and contralateral delays, Δ*T*_*inhi*_ and Δ*T*_*inhc*_, proposed in Myoga et al. (as described in the Introduction). We therefore tuned the delays to a starting scenario in which all four inputs to the MSO PCs arrive simultaneously when Δ*T*_*inhi*_ = Δ*T*_*inhc*_ = 0. As the recorded time for additional intermediate synaptic contact between GBCs and MNTB PCs or LNTB PCs was equal to 0.11 ms, we obtained the values displayed in Table III. Since no information was available about the exact values of the inter-population connections, all the other synaptic delays were kept to the default value of 1 ms.

**TABLE III.** Timings of connections between populations.

| Source pop | Dest pop | Delay (ms) |
| --- | --- | --- |
| GBCs | MNTBC | 0.45 |
| GBCs | LNTBC | 0.45 |
| SBCs | MSO (ipsilateral) | 2 |
| SBCs | MSO (contralateral) | 2 |
| LNTB | MSO | $1.44 + \Delta T_{inhi}$ |
| MNTB | MSO | $1.44 + \Delta T_{inhc}$ |

Finally, in Table 1 of *Supplementary*, we show the weights used for all synaptic connections in the network.

### C. Rate Analysis

Once studying the rates of the modeled auditory nuclei under different sound stimuli, we noticed how inner-population behavior often varied depending on the CF of their cells. We then analyzed the rate activity of clusters of *N* neurons centered around one having a specified CF. We call this particular subset a *CF cluster*. To observe the overall behavior of entire populations or *CF clusters* under different input conditions, we computed the average spiking rate per neuron. This metric counts all spikes recorded in a simulation from the analyzed population or *CF cluster* and divides it by its neuron count. This shows average activity changes depending on azimuthal angle. To instead observe the activity differences of different *CF clusters* within the same neural population, we used *rate matrices*: as heatmaps, they show the average spiking rate for a single neuron of different *CF clusters*, arranged on the y-axis, according to different azimuth angles of input sound, on the x-axis. Finally, for the analysis of LSO and MSO behavior, we computed the difference between the rate matrices of the left and right nuclei and normalized each of these *difference matrices* by the maximum difference across all *CF clusters* and azimuth angles. We present a graphical overview of this computation stage in Figure 1 of *Supplementary*.

**Fig. 1.**
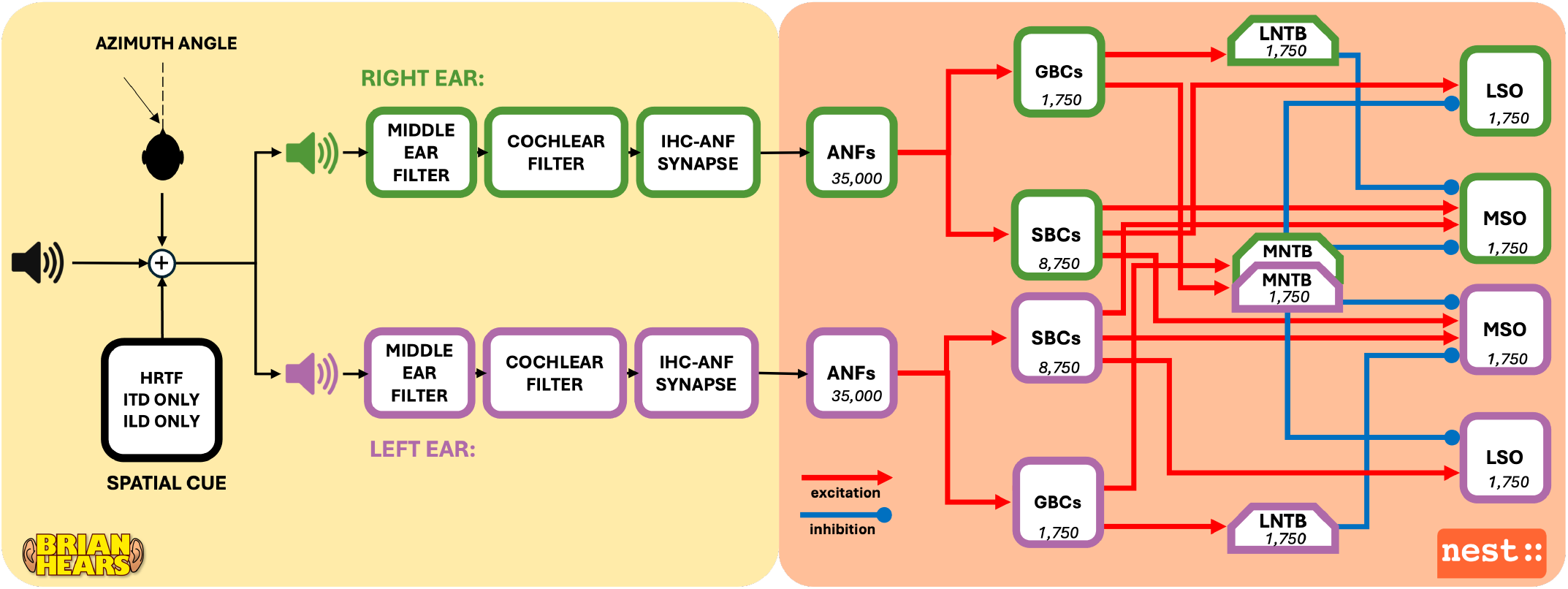
The end-to-end diagram of the proposed model, made of a peripheral section (yellow area), from sound to the spiking output of auditory nerve fibers (ANFs), implemented in Brian2Hears [35] and a brainstem section (orange area), developed in NEST [36]. Numbers inside the blocks indicate the neurons in each neural population. Red and blue arrows indicate excitatory and inhibitory synaptic connections, respectively.

### D. Simulator

For faster simulation times, we used a remote server with the following technical specifications: (1) Intel(R) Xeon(R) CPU E5-2690 v3 @ 2.60GHz (2) 16 CPUs (3) 1 thread per core (4) 64 GB RAM memory The server was equipped with Ubuntu 20.04.5 LTS.

## III. Results

The developed realistic computational model produced results that show both validating and novel features. In this section, we focus on novel results, but we first highlight the following as validating behavior shown by our model: (1)ANFs for different sounds show tuning properties consistent with literature behavior – tonotopic tuning, phase-locking, onset behavior (Figure 2 of *Supplementary*); (2) ANFs lose the capability to phase lock for pure tones with frequencies higher than 1.5 kHz (Figure 3 of *Supplementary*); (3) ANF spiking behavior changes consistently according to azimuth angle, with onset and firing rates reflecting differing ITDs and ILDs (Figure 4 of *Supplementary*); (4) bushy cells (SBCs and GBCs), targeted by ANFs, show improved phase-locking, enabling time-enhanced signals to reach MSO and LSO (Figure 5 of *Supplementary*).

**Fig. 2.**
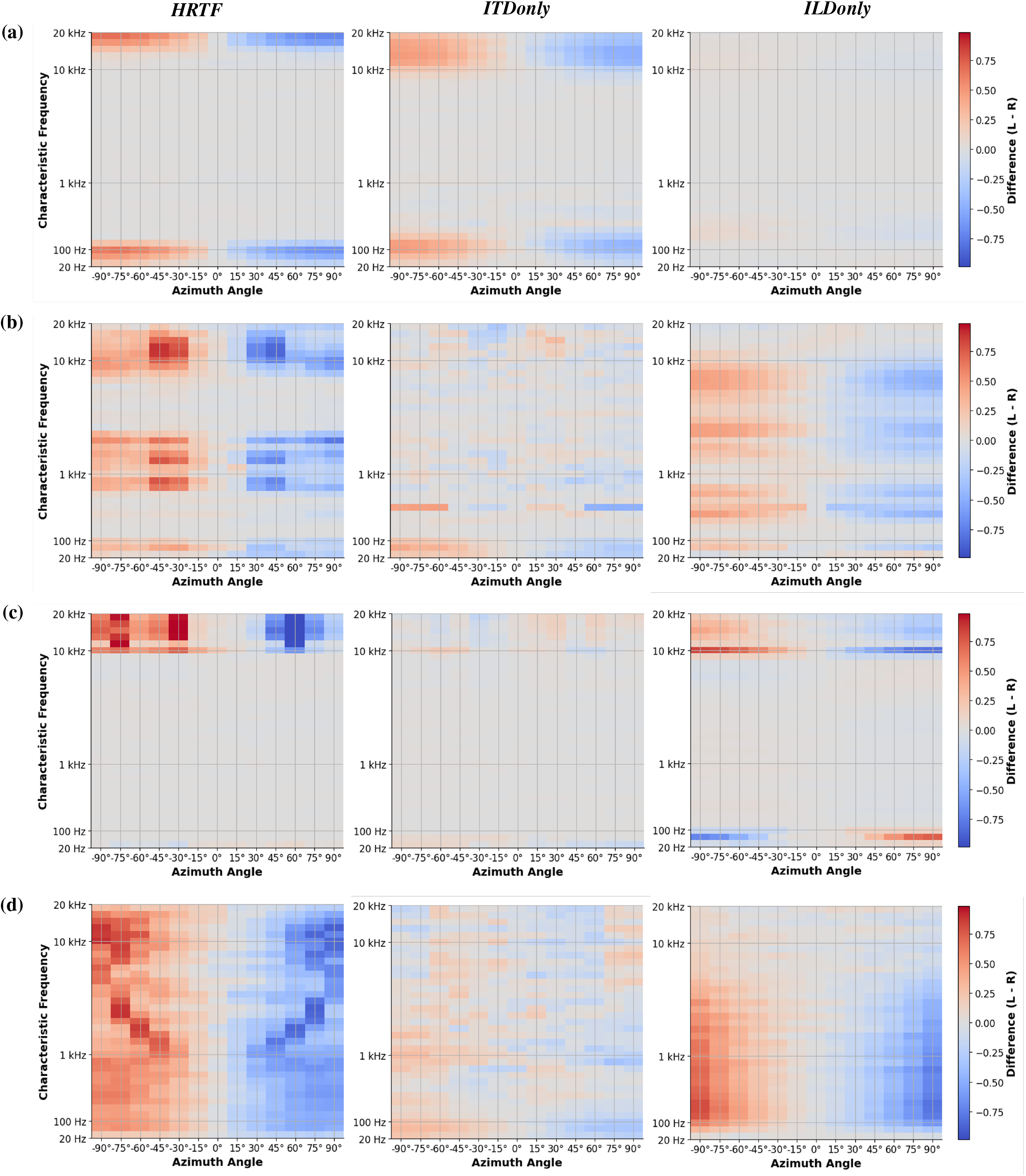
LSO’s difference matrices for 0.1 kHz (a), 1 kHz (b), 10 kHz (c) tones and white noise (d) under the three cue configurations.

**Fig. 3.**
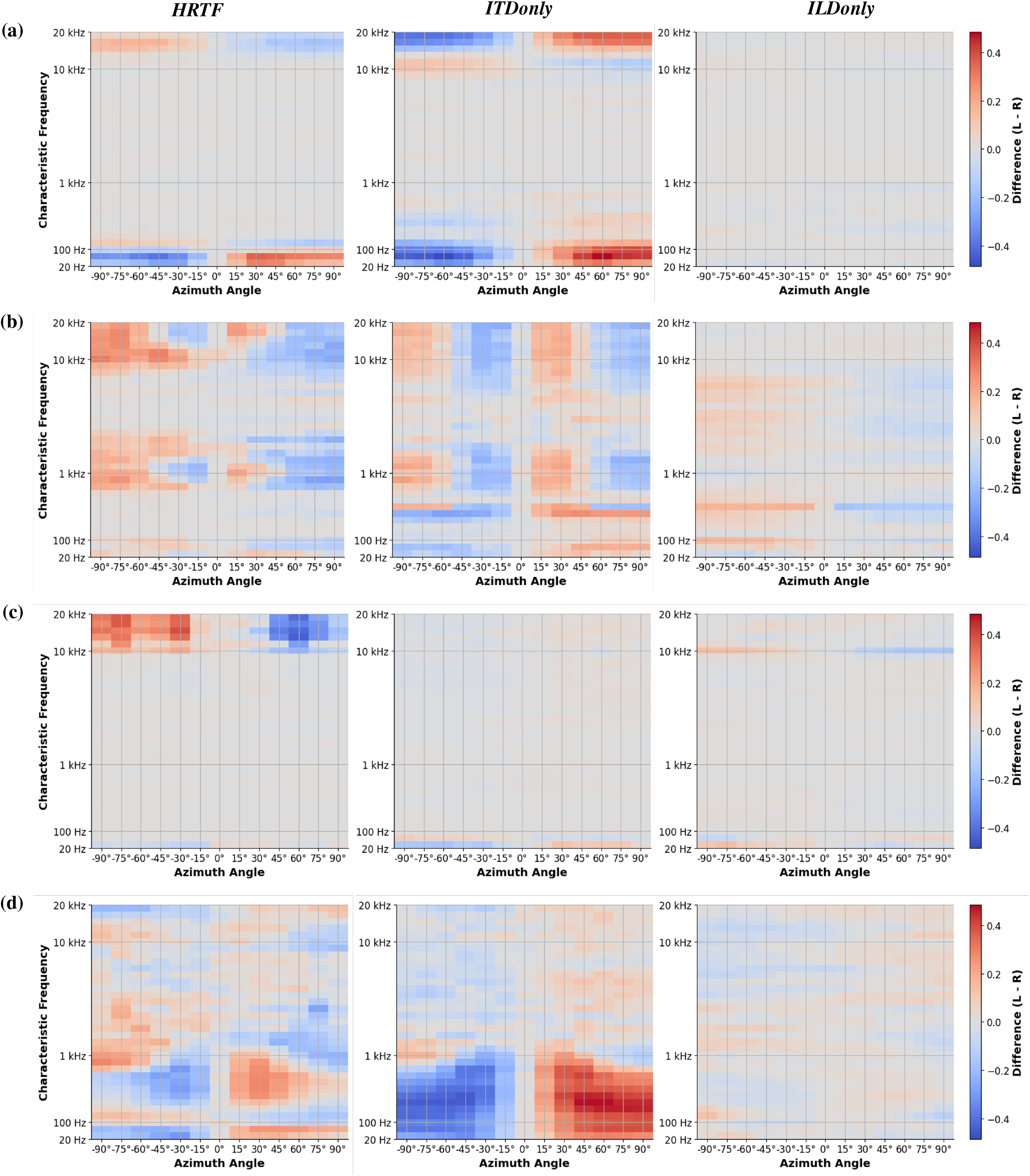
MSO’s difference matrices for 0.1 kHz (a), 1 kHz (b), 10 kHz (c) tones and white noise (d) under the three cue configurations.

**Fig. 4.**
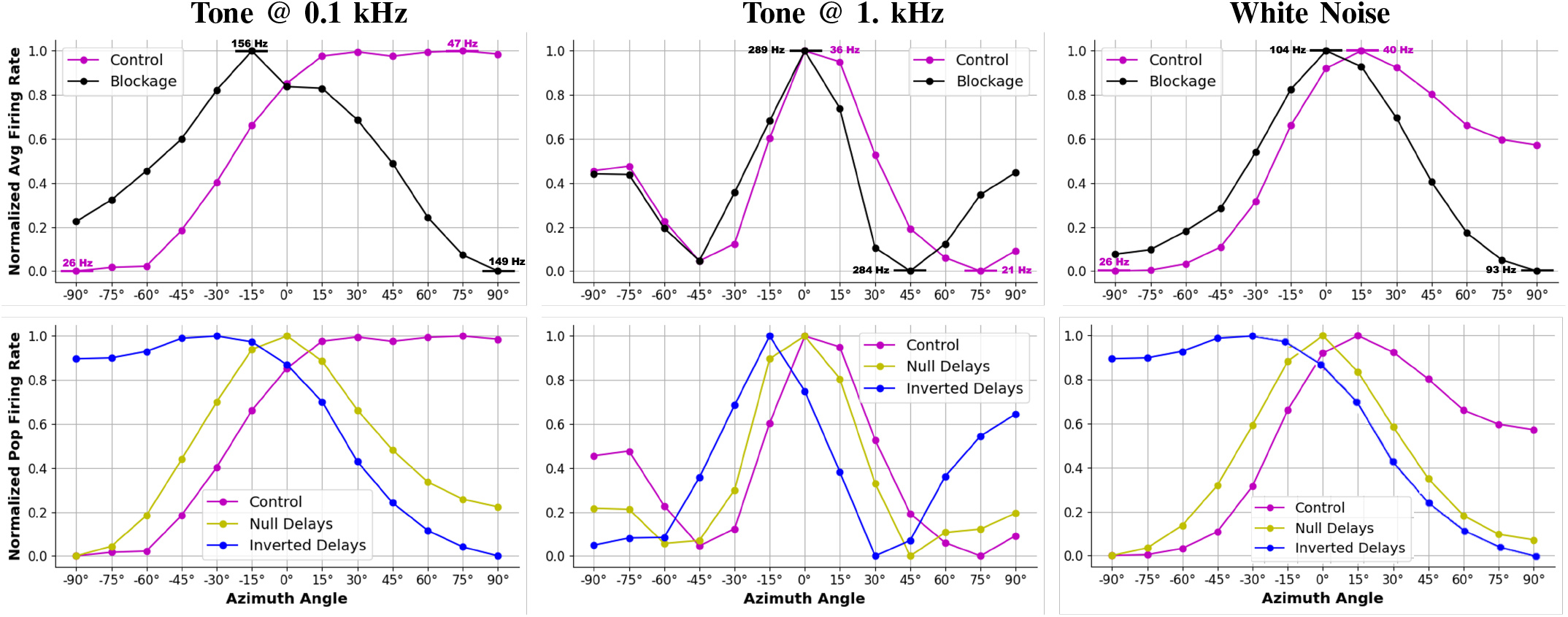
MSO activity for 0.1 kHz (left), 1 kHz tones (center), and white noise (right): above, complete inhibition blockage increased the overall firing rate and shifted the peak in MSO response towards 0°; below, varying combinations of Δ*T*_*inhi*_, Δ*T*_*inhc*_ moved the peak of MSO response towards more ipsilateral locations.

For our investigation, we focused on the open problems highlighted in the Introduction (Section I):

A. **Duplex theory.** When stimulated by pure tones, neural activity in MSO and LSO confirms psychoacoustic results for ITD and ILD frequency dependence [21].
B. **Presumed cue segregation between LSO and MSO.** Simplistic views of complete segregation between LSO and MSO roles do not hold when investigating activity patterns for tones at varying frequency [18], [22]. Stimulation by white noise points toward an amplified segregation when compared to tones.
C. **Inhibition role in MSO ITD-coding.** Earlier results with *in vivo* inhibition blockage [14] were confirmed by our *in silico* results, increasing evidence for inhibition’s relevance.
D. **Whole-network effects of inhibition timing.** Earlier results on inhibition timing influence on peak firing, conducted in brain slices [17], still stand when the same principles are applied at a network level.

### A. Duplex theory

Psychoacoustic results point to worsened localization accuracy at frequencies where both ITD and ILD have a limited effect, explained by considering these two cues as the main drivers for solving azimuth sound localization. The important role of ITD and ILD is similarly evidenced by the neural activity in MSO and LSO, where the neural circuit for azimuthal sound localization points to. ITD is the main driver for low-frequency responses, and as input frequency rises, it progressively loses its relevance in favor of ILD. This holds true for both LSO and MSO, even if their rate response is different. Across different *CF clusters*, LSO neurons uniformly exhibit a sigmoid-like firing-rate function of sound azimuth with ipsilateral preference, consistently with literature [18]. In accordance with duplex theory, for a pure tone at 0.1 kHz (Figure 2a), the spatial tuning exclusively derives from ITDs, since ILD contribution is practically negligible. For tone at 1 kHz (Figure 2b), all the fibers increasingly rely on ILDs, and finally, with tones at 10 kHz (Figure 2c), LSO neurons respond solely to ILDs, preserving the sigmoid form but devoid of ITD influence.

Different from the LSO, MSO’s *CF clusters* tuning is heterogeneous along stimulus conditions. For tones at 0.1 kHz (Figure 3a), low-CF neurons displayed a sigmoid curve with contralateral preference in the HRTF scenario, in accordance with literature; high-CF fibers instead exhibited LSO-like responses, showing a sigmoid behavior with ipsilateral preference. Nevertheless, when stimuli were restricted to ITD cues only, LSO-like behavior for high-CF fibers disappeared, showing homogeneous MSO sensitivity to ITDs across all the population. Again, duplex theory was confirmed, since ILD response was null for tone at 0.1 kHz. For the 1 kHz tone (Figure 3b), we observe a complex pattern in the MSO matrices: when subjected only to ITDs, the low-CF neurons retain sigmoid with contralateral preference curves; moving toward medium- and high-CF, units curves start to change, adopting a contralateral-peaked shaped, described in the matrix by a red zone (left MSO stronger than the right one) alternated to a blue zone (peak of the right MSO, stronger than the left) for azimuth angles between −90° and 0° (coming from the left side), then a red zone (peak of the left MSO, stronger than the right) and finally a blue zone (right MSO stronger than the left one). In the HRTF scenario, this behavior is still present, but it is less evident, potentially disturbed by the LSO-like (so sigmoid with ipsilateral preference) behavior enforced by ILDs. At 10 kHz (Figure 3c), contralateral preference responses vanish: high-CF units respond exclusively to ILD via LSO-like sigmoids with ipsilateral preference. In both nuclei, the 3×3 *difference matrices* associated with pure tones exhibit spatial tuning (i.e., non-gray areas) only in regions that align with *CF clusters* where phase-locking activity is present in ANFs (see Figure 2 in *Supplementary*).

### B. Presumed segregation between LSO and MSO

Literature frequently proposes a conventional division of tasks for MSO and LSO concerning the processing of ITDs and ILDs. Nevertheless, our results for pure tone stimulation show activity to be driven by ITDs for 0.1 kHz tone and by ILDs for 1 and 10 kHz tones in both nuclei. This confirms a dependence on tone frequency as stated by the duplex theory, more than a strict segregation between the two nuclei. However, in response to white noise stimulation, a functional division between LSO and MSO seems to emerge. LSO units respond again with a sigmoidal firing-rate function with ipsilateral preference (Figure 2d), generated mostly by ILD cues across all *CF clusters*, with only a small ITD contribution. For the MSO (Figure 3d), we observe, as for the 1 kHz tone, a sigmoid function with contralateral preference for low-CF neurons, which becomes peaked for medium to high CF clusters. Again, this behavior is refined when MSO is subjected to the *ITDonly* scenario, whereas no relevant tuning is detected when subjected to ILD only ones, even at high *CF clusters*.

### C. Inhibition role in MSO ITD coding

To understand the role of inhibition in computing ITDs, we performed a set of simulations focusing on the *ITDonly* cue configuration applied to tones at 0.1 and 1 kHz, as well as to white noise. We chose these three sounds because they were the ones that elicited the most evident contralateral preference, whereas the 10 kHz tone produced an ipsilateral preference. To assess the global role of inhibitory inputs to the MSO, we set their weights to 0, replicating *in silico* the strychnine-induced inhibition blockade applied to MSO neurons *in vitro* in [14]. Removing inhibition resulted in a higher firing rate across azimuth angles compared to the control simulations (Figure 4). To compare the tuning curve shapes, as done in [14], we performed a min-max normalization; we included the absolute values of minimum and maximum firing rate in the figures. For all sounds, inhibition removal led to a complete loss of contralateral preference, with the peak response shifting toward 0°.

### D. Whole-network effects caused by inhibition timing

Our last investigation underpins how we structured the MSO model: in [17], the authors demonstrated that in gerbil brain slices, inhibitory inputs shift the timing of peak firing rate; specifically, the magnitude of this shift depends on the arrival time of inhibitory inputs relative to their corresponding excitation. By exploring the input space, the timing of optimal coincidence detection was found. For the previous results, we strictly adopted this optimal input: Δ*T*_*inhi*_ = 0.2*ms*, Δ*T*_*inhc*_ = 0.4*ms*. Nevertheless, by experimenting with varying Δ*T*_*inhi*_ and Δ*T*_*inhi*_, we demonstrate that relative arrival time is similarly able to shift peak activity *population* rates. In Figure 4 we show how activity varies in two scenarios with respect to the optimal: without any relative arrival delay (Δ*T*_*inhi*_ = Δ*T*_*inhc*_ = 0*ms*) and with timing values shifted to the opposite end of the range (Δ*T*_*inhi*_ = 0.4*ms*, Δ*T*_*inhc*_ = 0.2*ms*). We found that removing relative arrival delay caused the MSO activity range to peak at 0°, while using values opposite to the optimal lead to a symmetric mirroring of the optimal curve, showing now an ipsilateral peak.

## IV. Discussion and conclusions

Our computational investigation aimed to address key unresolved questions regarding azimuthal sound localization in the human superior olivary complex, focusing on ILD and ITD processing in the MSO and LSO. By flexibly varying sound stimuli, configuring different cue conditions, and independently analyzing distinct *CF clusters* in the MSO and LSO, thus accounting for the tonotopic organization of neuronal populations, we were able to provide consistent and quantitative answers to several ongoing debates about the neural mechanisms underlying this task. First, we established a robust platform to evaluate the relevance of the duplex theory [2] in MSO and LSO neural activity. Our findings support the classic division of roles, where ITDs and ILDs are preferentially used to encode the location of pure tone stimuli depending on frequency, a division long supported by psychoacoustic and behavioral evidence but never validated experimentally, especially in humans. Consistent with this, we did not observe a strict anatomical segregation between MSO and LSO for ITD and ILD coding of pure tones. However, this functional dichotomy emerged when the network was presented with more complex stimuli such as white noise, suggesting cue-dependent specialization between the two nuclei. We further used our model to address one of the most debated questions in sound localization: how ITDs, which lie in the microsecond range, are coded by neuronal activity in the MSO. Our results support the hypothesis proposed by Brand et al. [13] and later investigated by Pecka et al. [14], confirming the role of inhibition in shifting MSO rate activity toward contralateral angles. Our findings argue against the hypothesis that this shift is an off-target effect of strychnine (used to block glycinergic inhibition in [14]), as suggested in [9], and instead reinforce the key role of inhibition itself. We also investigated the influence of relative arrival times between ipsilateral inhibition and excitation on MSO activity, translating the optimal timing values identified by Myoga et al. [17] through conductance-clamp experiments in gerbil brain slices. Our simulations demonstrate that fine-tuned timing between these inputs is a crucial determinant of the contralateral shift in ITD sensitivity. Moreover, we have verified that the obtained results are not limited to the specific HRTF taken into account. In fact, we repeated the same analysis for the four sounds studied with 5 HRTFs recorded from different subjects. We therefore verified that the results are not dependent on the subject and can be generalized to a wider range of subjects or conditions (see Figure 8 and 9 of *Supplementary*). While our network offers valuable insights, several important limitations must be acknowledged. Most notably, the model lacks spectral cue processing in the ventral cochlear nucleus, which plays a vital role in resolving front-back ambiguities and vertical localization. This omission restricts our ability to fully characterize three-dimensional spatial hearing. Additionally, our stimulus set was limited to pure tones and white noise. Given that white noise evoked a clearer functional segregation between MSO and LSO, it will be important to evaluate the outcome for more complex stimuli, such as clicks or speech, in future work. Furthermore, our model does not include higher-order auditory centers such as the inferior colliculus, the thalamic medial geniculate body, or the primary auditory cortex. As a result, we cannot assess how binaural information is refined, integrated, or modulated at later stages, nor can we explore feedback and top-down influences on auditory processing. Lastly, the current model lacks plasticity mechanisms that are known to play essential roles in adaptive sound localization, particularly during development and following auditory experience. The precise timing between inhibitory and excitatory inputs, which we showed to be central for ITD coding, may itself be shaped through plastic processes, an important avenue for future research. Beyond these constraints, the modeling framework developed in this study serves as a comprehensive and flexible testbed for investigating the responses of auditory brainstem nuclei to spatially distributed acoustic stimuli. By capturing key mechanisms underlying ILD and ITD processing and allowing for controlled manipulation of input cues and neuronal properties, our model provides a solid foundation for future studies on binaural processing and auditory spatial encoding. To facilitate further development and experimentation, we provide full access to the network code here: https://anonymous.4open.science/r/sound_localization_model-DE63

