## Supplementary File for "A human spiking computational model to explore sound localization"

### Supplementary Materials

TABLE I
SYNAPTIC WEIGHTS BETWEEN POPULATIONS

| Presynaptic population | Postsynaptic population | Synaptic Weight (nS) |
|---|---|---|
| ANFs | SBCs | 35.0 |
| ANFs | GBCs | 7.0 |
| GBCs | LNTBCs | 20.0 |
| GBCs | MNTBCs | 30.0 |
| SBCs | LSO | 10.0 |
| MNTBCs | LSO | -10.0 |
| SBCs | MSO | 9.0 |
| MNTBCs | MSO | -40.0 |
| LNTBCs | MSO | -40.0 |

#### I. Furter Analysis

*1) Rasterplots and PSTHs:* We employed rasterplots to qualitatively assess the behaviors of ANFs and SBCs in response to various input sounds. To differentiate between the main observable behaviors within diverse *CF clusters*, rasterplots display the CF scale on the y-axis, mapping each analyzed population (including those beyond ANFs) with the ERB scale [1]. When it was necessary to compare different populations, such as between left and right nuclei or across different azimuth angles, a more condensed visualization became essential. Thus, we adopted Post-Stimulus-Time-Histograms, which count the absolute number of spikes within time bins that are 1 ms wide.

*2) Vector Strength:* To have a more quantitative measure of the phase-locking ability, we used vector strength (R), a measure of phase synchrony, or how well spike timings are synchronized to a specific phase of a periodic stimuli (like sound waves). Because it directly relates spike timing to the phase of an input frequency, phase-locking analysis only make sense when applied to tonal sounds. We adapted a formula from [2] to capture this. First, spike times are converted into phases within the cycle of the stimulus using:

$$\textit{phase} = 2\pi \times (\textit{spike time} \bmod \frac{1}{\textit{frequency}}) \times \textit{frequency} \quad (1)$$

This maps each spike time to an angular value on the unit circle relative to the sound's frequency. The vector strength is then calculated as the magnitude of the mean resultant vector from these phases, using

$$R = \sqrt{(\text{mean}(\cos(\textit{phase})))^2 + (\text{mean}(\sin(\textit{phase})))^2} \quad (2)$$

A value close to 1 indicates that spikes are tightly clustered at a particular phase, reflecting strong phase-locking and precise temporal coding of that frequency, while a value near 0 indicates a lack of synchronization. Thus, vector strength provides a clear and quantitative measure of how well neural activity aligns with the frequency of an auditory stimulus. Measuring R across the frequency spectrum is relevant to investigate how it changes depending on the distance from the cell characteristic frequency.

#### II. Further Results

##### A. Auditory Nerve Fibers (ANFs)

The following images are purposed to validate ANFs behavior with respct to their biological counterparts.

In Figure 2, we illustrate the spiking activity of ANFs in response to our set of sounds, all presented from a 0° azimuth angle. The combined effects of HRTF filtering and cochlear processing introduced a delay of approximately 7 ms from sound onset to the point where ANFs began responding. Additionally, the responses varied depending on the characteristic frequency (CF) of the ANFs and can be categorized into four distinct types: *tuning to their CF* (cells firing tonotopically at their own CF), *phase-locking* (cells firing at input tone frequency), *no tuning* (spiking dominated by spontaneous activity), and *onset activation* (a single spike at stimulus onset followed by a *no tuning* state). For tones at 0.1 kHz and 1 kHz, ANFs with low and medium CFs (20 Hz–2 kHz) showed *tuning to their CFs*. Particularly evident for the 1 kHz tone within a *CF cluster* centered around 1 kHz. In contrast, high-CF ANFs (10–20 kHz) exhibited *phase-locking* for both tones. Between these regions, ANFs responded with *no tuning* to the 0.1 kHz tone and with *onset activation* to the 1 kHz tone. In the case of a 10 kHz tone, low and medium CFs yielded *no tuning*; medium-to-high CFs showed *onset activation*; and high CFs again exhibited *phase-locking*. As expected, the white noise stimulus evoked *tuning to the respective CFs* across all modeled ANFs. To asses the tonotopic arrangement of ANFs, we computed vector strength in response to tones at 0.1, 0.5, 1, and 1.5 kHz for low-CF ANFs (20-2000 Hz). The results (see Figure 3a) show that vector strength (R) peaked in ANFs whose CF matched the input frequency, that is, within the relevant *CF cluster*. Notably, the width of this cluster (i.e., the range of CFs with high R values) increased with the frequency of the stimulus. Figure 3b illustrates how four ANFs with CFs of 100, 500, 1000, and 1500 Hz responded to multiple tones in terms of vector strength. ANFs displayed higher R values for tones near their CF and gradually lost tuning for more distant frequencies. Azimuthal variation is critical for sound localization, especially for binaural cues, and can be inferred from differences in activity between left and right populations. Figure 4 presents ten post-stimulus time histograms (PSTHs) showing ANFs activity for two input sounds originating from five azimuth angles: $-90°$, $-45°$, $0°$, $+45°$, and $+90°$, covering the frontal azimuthal space from left to right in $45°$ steps. For both sounds, we observed coherent differential activity patterns across hemispheres. At $-90°$, left ANFs exhibited earlier and stronger responses; this dominance diminished at $-45°$ and became symmetric at $0°$. At $+45°$ and $+90°$, right ANFs led in both timing and strength. This pattern of greater responsiveness for the ipsilateral population was particularly pronounced for the white noise stimulus.

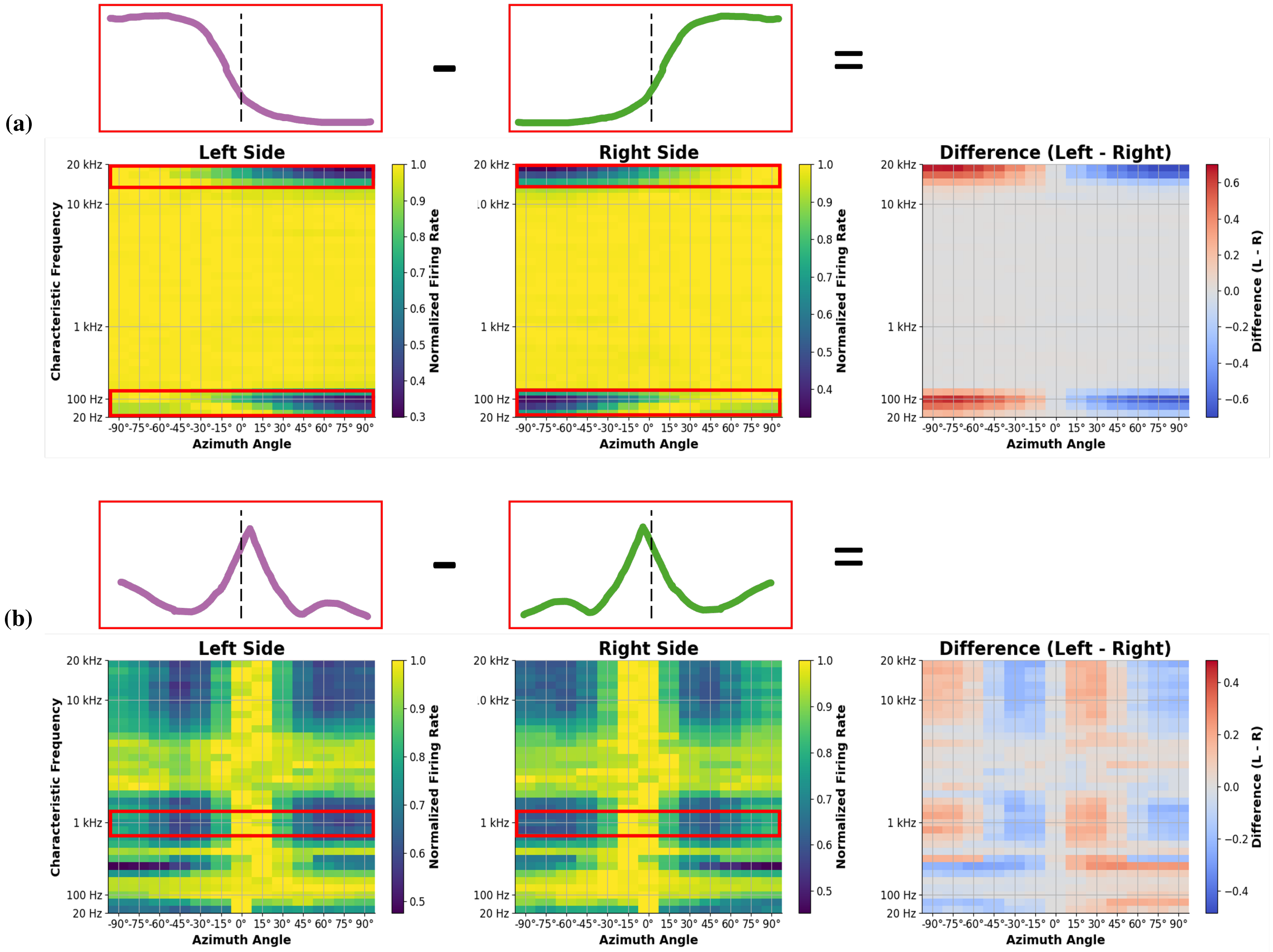


Fig. 1. Computation scheme of *difference rate matrices* used to analyze LSO and MSO responses to various sound stimuli and cue conditions:
(a) LSO response to 0.1 kHz tone under *HRTF* cue condition: sigmoidal with ipsilateral preference rate activity is observable at low and high CF fibers.
(b) MSO response to 1.0 kHz tone under *ITDonly* cue condition: peaked with contralateral preference rate activity is observable at 1 kHz CF fibers.
Red boxes underline the correspondence of the obtained curves with the traditional LSO and MSO's rate responses found in literature

##### *B. Spherical and Globular Bushy Cells (SBCs and GBCs)*

Figure 5 demonstrates how phase-locking is refined at the bushy cell level in our network, in a manner realistic to their biological counterparts. We stimulated the system with a pure tone at 0.3 kHz. Raster plots show that the qualitative response pattern across CFs remained consistent across populations, but tuning became sharper in SBCs and GBCs. We quantified this effect within a CF cluster centered at 300 Hz and observed an increase in R from ANFs to SBCs and GBCs. Beyond input sound type and CF, azimuth angle is the third major factor influencing the input to our network.

##### *C. Lateral and Medial Superior Olives (LSO and MSO)*

Figures 6 and 9 display the *average firing rate* of LSO and MSO neurons under the four different sound stimuli and three cue conditions analyzed in the main text. LSO neurons were studied globally, since rate activity across population with different CF was homogeneous. MSO's activity across different CF cluser was instead highly heterogeneous; thus, in Figure 9 we chose to display for pure tones stimulations just the CF cluster centered on the pure tone frequency. For the white noise, the average firing rate was computed globally, by considering the entire MSO population, analogously at what was been done for the LSO.

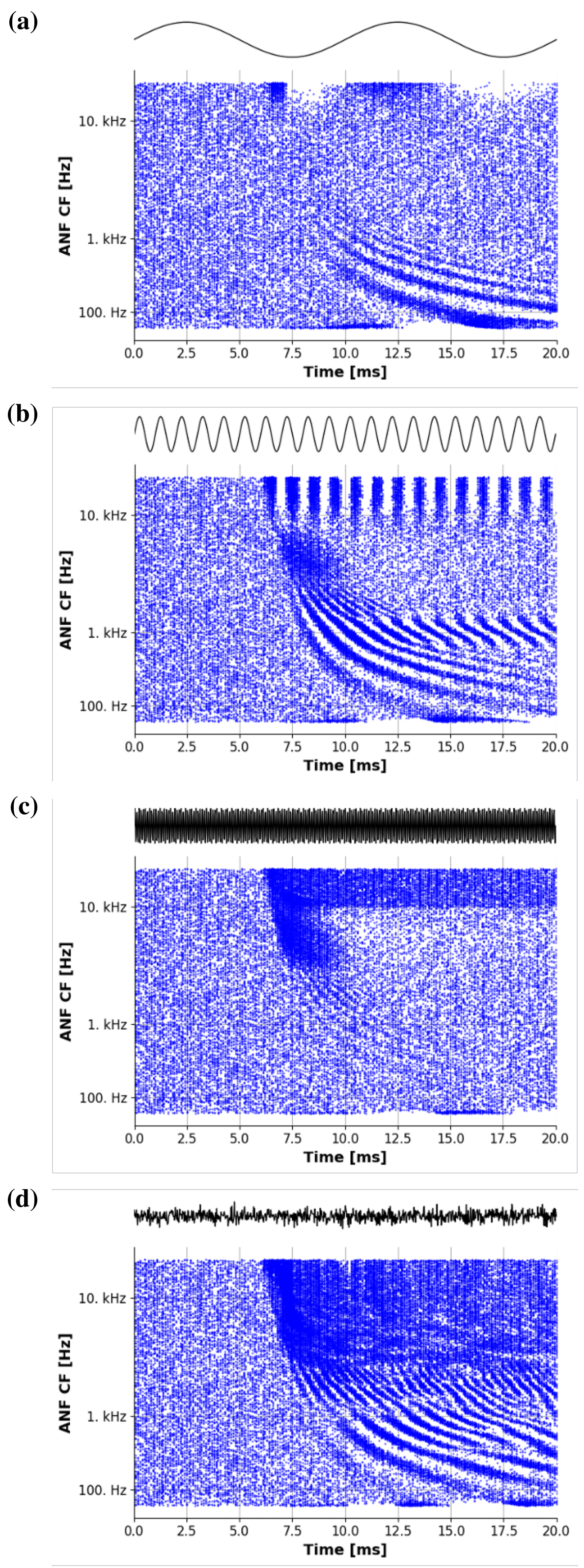


Fig. 2. ANFs tonotopic response to 0.1 kHz (a), 1 kHz (b), 10 kHz (c) tones and to white noise (d).

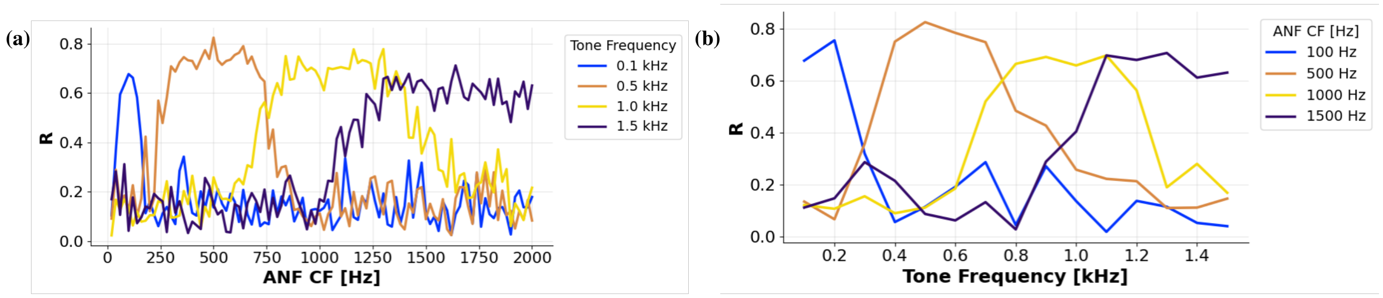


Fig. 3. Phase-locking of ANFs measured by computing the vector strength $R$. (a) $R$ measured among all ANFs with CF between 20–2000 Hz for four different tones of frequency 0.1, 0.5, 1, and 1.5 kHz. (b) $R$ measured among different tones for four different ANFs with CF equal to 100, 500, 1000, and 1500 Hz.

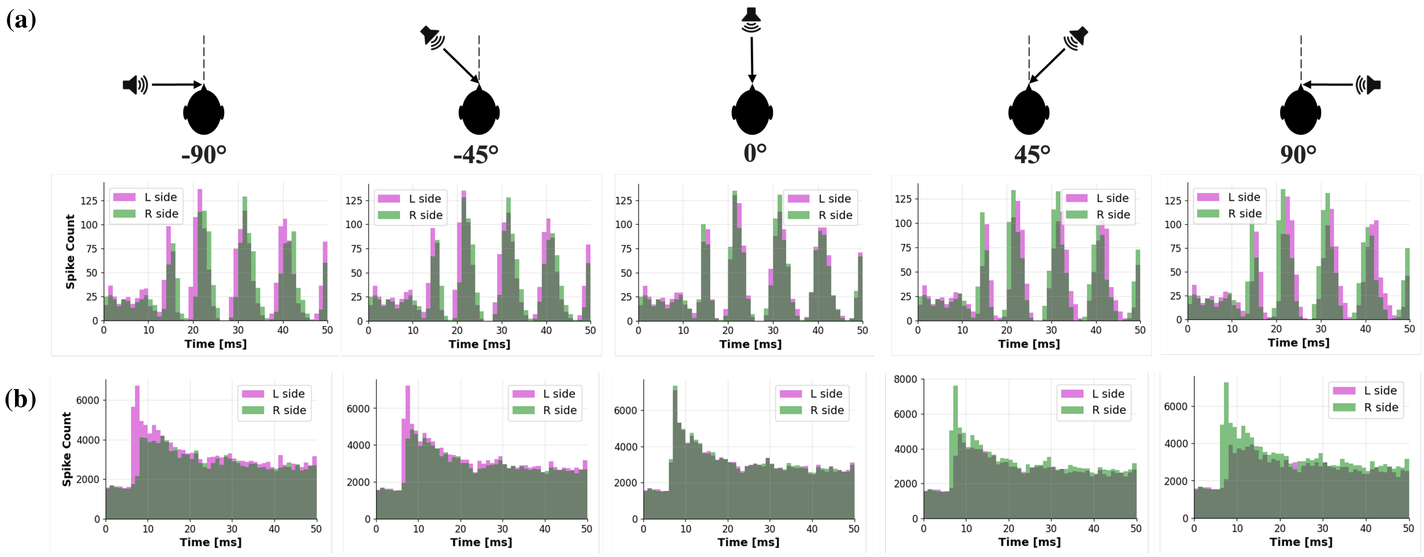


Fig. 4. PSTHs for the first 20 ms of ANFs response to 0.1 kHz tone (a) and white noise (b) coming from five different azimuth locations, spanning all the frontal horizontal space

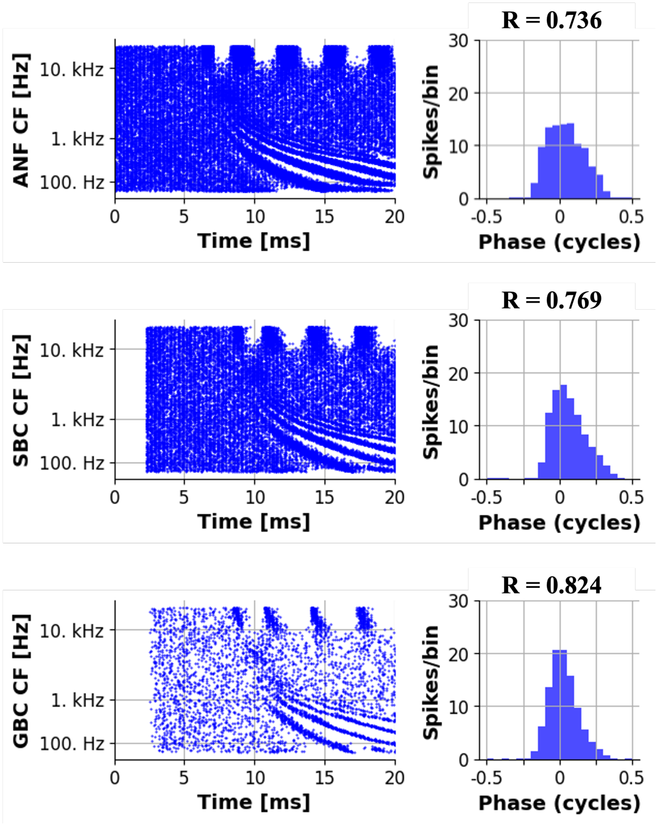


Fig. 5. Phase-locking enhancement in SBCs and GBCs due to the convergence of many ANFs. The phenomenon is observable qualitatively from the population rasterplots (in response to a 0.3 kHz tone, replicating experimental setup in [3]) and is quantified from the computation of vector strength (R) values for ANF, SBC and GBC with CF = 300 Hz (so same as input tone), which can also be visualized in the vector strength plots

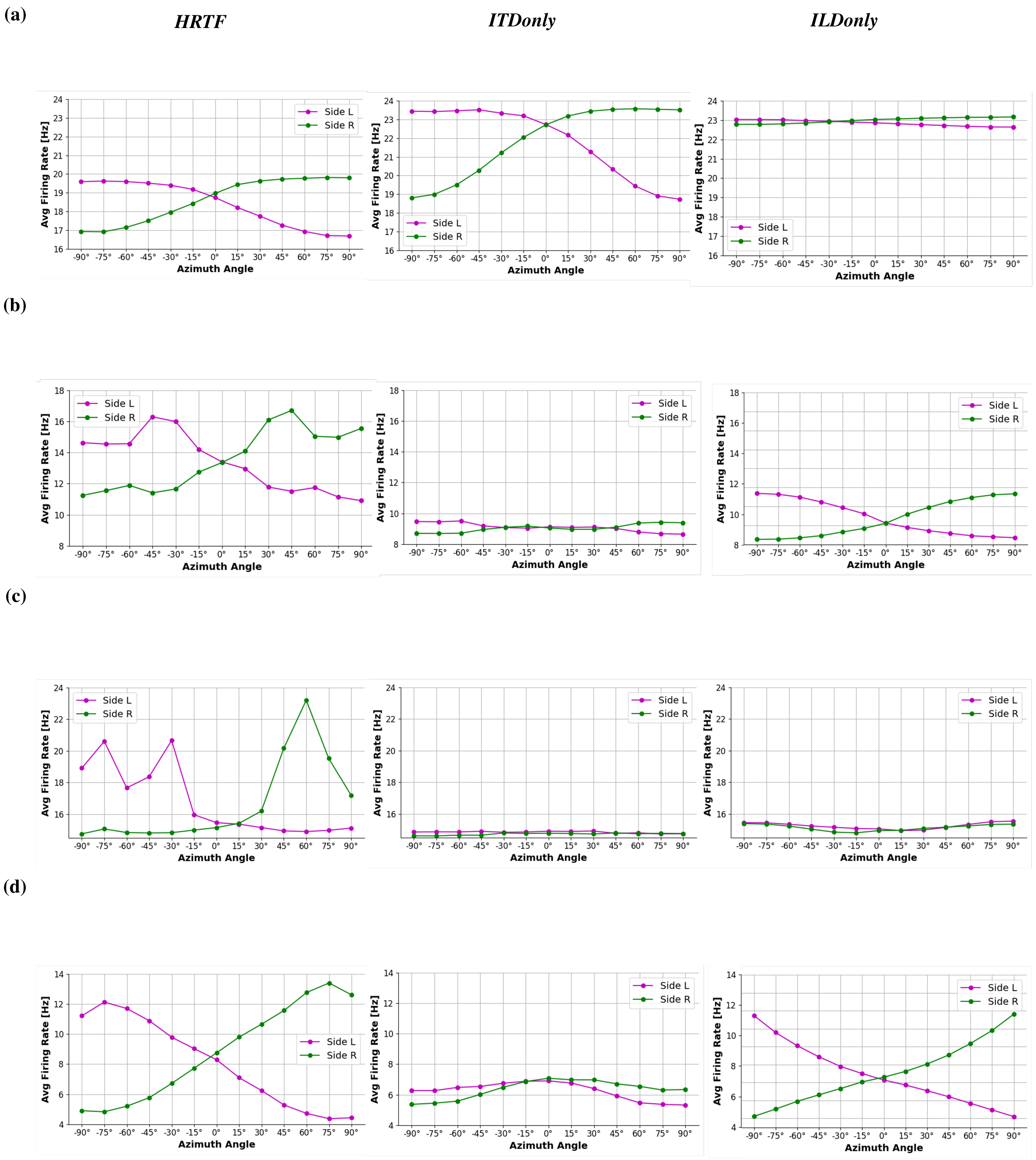


Fig. 6. LSO's difference matrices for 0.1 kHz (a), 1 kHz (b), 10 kHz (c) tones and white noise (d) under the three cue configurations. Average firing rate is computed considering the contribution of the entire LSO population (CF between 20 Hz and 20 kHz)

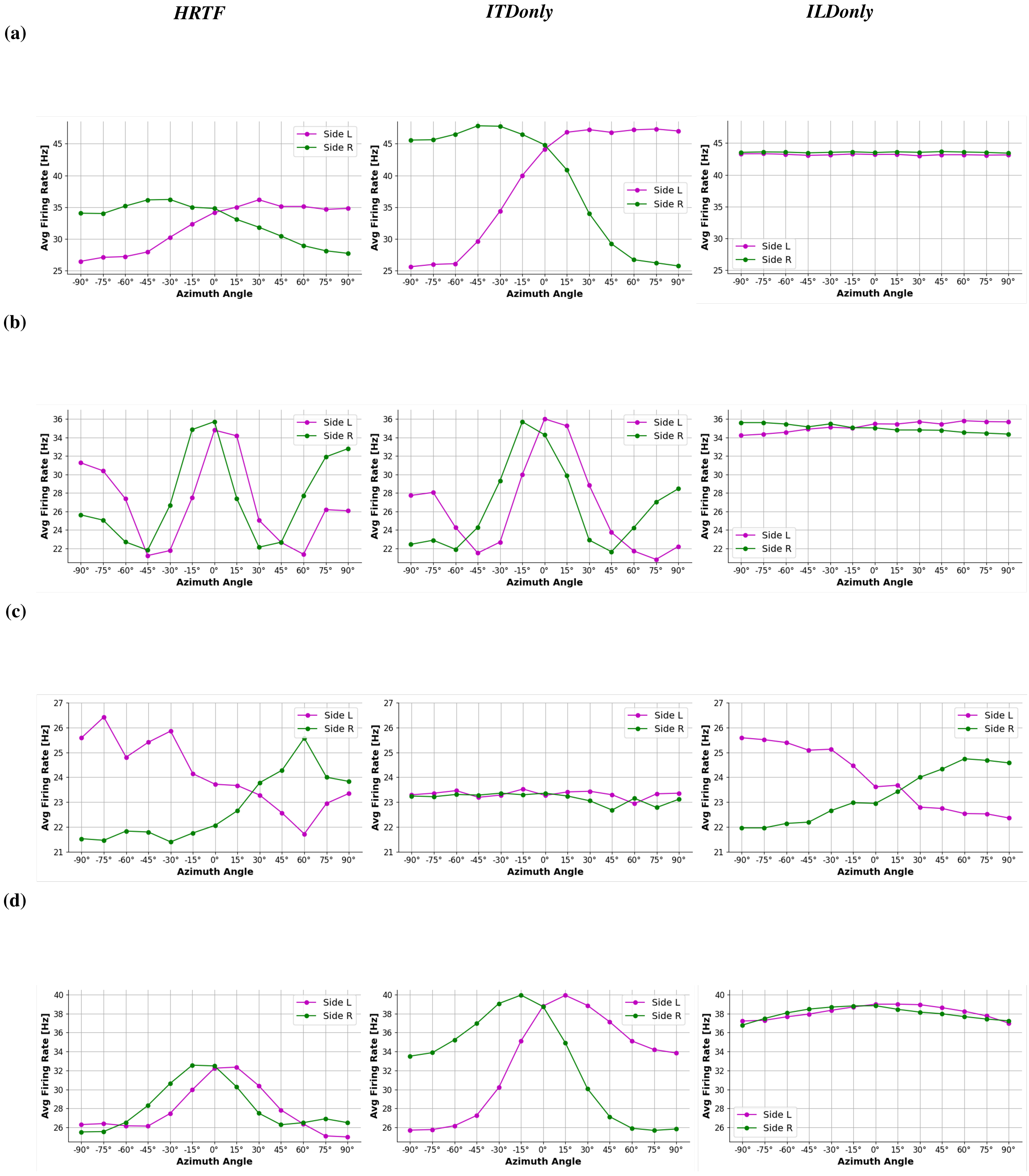


Fig. 7. MSO's difference matrices for 0.1 kHz (a), 1 kHz (b), 10 kHz (c) tones and white noise (d) under the three cue configurations. (a,b,c) Average firing rates are computed considering the contribution of a single CF cluster centered on the pure tone frequency, containing 50 neurons. (d) Average firing rate is computed considering the contribution of the entire MSO population (CF between 20 Hz and 20 kHz)

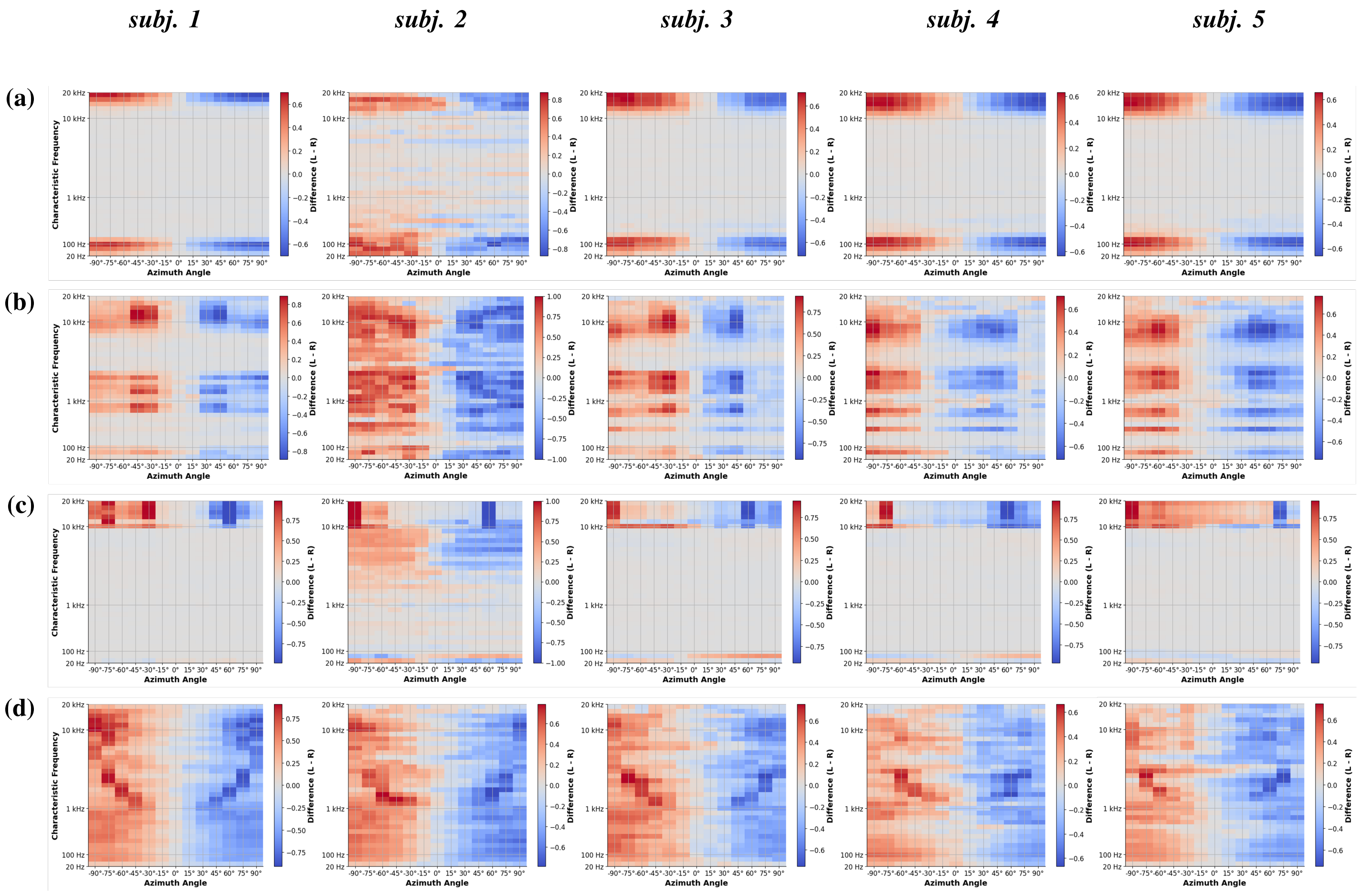


Fig. 8. LSO's difference matrices for 0.1 kHz (a), 1 kHz (b), 10 kHz (c) tones and white noise (d) for the five different HRTFs analyzed, corresponding to five different subjects in the IRCAM database [4]. The nucleus's activity proves to be consistent among the five subjects analyzed.

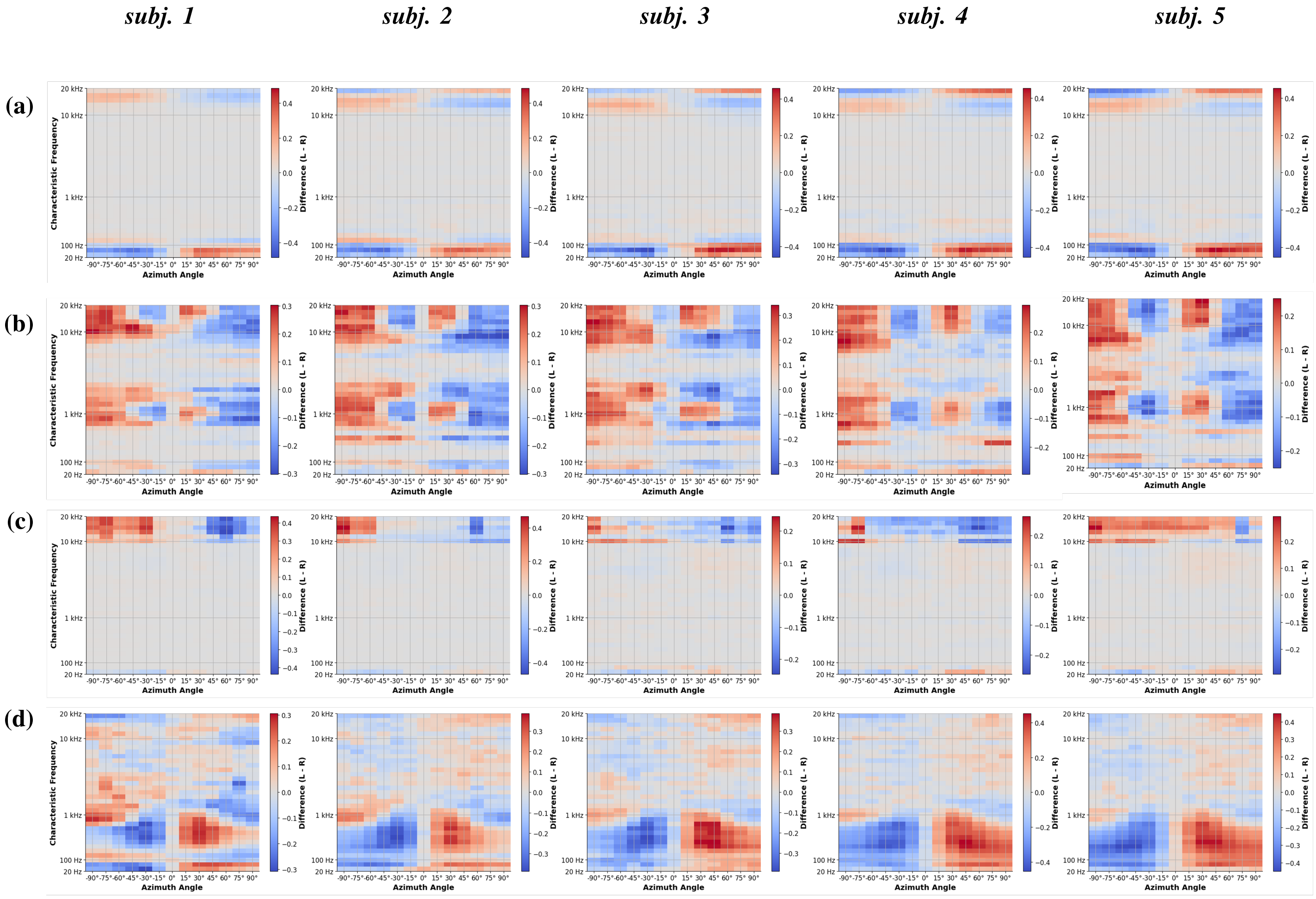


Fig. 9. MSO's difference matrices for 0.1 kHz (a), 1 kHz (b), 10 kHz (c) tones and white noise (d) for the five different HRTFs analyzed, corresponding to five different subjects in the IRCAM database [4]. The nucleus's activity proves to be consistent among the five subjects analyzed.
